# LIN-67 functionally interacts with heterochronic miRNAs and regulates developmental timing in *Caenorhabditis elegans*

**DOI:** 10.1101/2024.12.31.630947

**Authors:** Jeffrey C. Medley, Roberto Perales, Belén Gaete Humada, Katja Doerfel, Ariana Levine, Ganesh Panzade, Christopher M. Hammell, Anna Zinovyeva

**Affiliations:** Division of Biology, Kansas State University, Manhattan, KS 66502; Cold Spring Harbor Laboratory, One Bungtown road, Cold Spring Harbor, NY 11724; Stony Brook University, Stony Brook, NY 11790; Laboratory of Human Retrovirology and Immunoinformatics, Frederick National Laboratory for Cancer Research, Frederick, MD 21702

**Keywords:** *C. elegans*, heterochronic, microRNA, KH domain, development

## Abstract

Temporal regulation of gene expression is required for developmental transitions, including differentiation, proliferation, and morphogenesis. In the nematode *Caenorhabditis elegans*, heterochronic microRNAs (miRNAs) regulate the temporal expression of genes that promote animal development. The heterochronic miRNAs lin-4 and let-7 are required during different stages of larval development and are associated with the miRNA-specific Argonaute ALG-1. In this study, we have identified *lin-67* as a heterochronic gene that negatively regulates *lin-4*, *let-7,* and *alg-1*. Loss of *lin-67* function restores proper developmental timing and stage-specific gene expression to hypomorphic *lin-4* and *let-7* mutants. We found that loss of *lin-67* resulted in a reduced number of seam cells, defects in alae formation, precocious expression of an adult-specific gene reporter, and sterility. LIN-67 contains a K homology (KH) RNA-binding domain and is a homolog of the Sam68-like splicing factor KHDRBS2. We show that LIN-67 localizes to the nucleus throughout animal development and is enriched in nuclear foci. Mutating the KH domain of LIN-67 abolished the nuclear localization of LIN-67, suggesting that the localization of LIN-67 is likely dependent on RNA-binding activity. We show that LIN-67 negatively regulates *lin-4* miRNA levels and restores normal levels of *let-7* to *alg-1* mutants, which can, at least in part, explain how *lin-67* suppresses *alg-1*. Our data indicate that *lin-67* is a novel heterochronic gene that regulates developmental timing and miRNA-dependent gene regulation in *C. elegans*.

## Introduction

During animal development, spatiotemporal control of gene expression drives genetic programs that give rise to diverse cellular and tissular functions (Buccitelli and Selbach, 2020; Davidson, 2010). Normal developmental timing requires timely inactivation and activation of different genes to establish and coordinate key developmental transitions. Disruptions in developmental programs give rise to heterochrony and are associated with several human disorders (Wilson, 1988; Smith and Flodman, 2018). microRNAs (miRNA) are non-coding small RNAs that play a central role as developmental timing regulators by repressing target genes’ activity (Bartel, 2018; Shang et al., 2023). Mature miRNAs are loaded into Argonaute proteins to form the miRNA-induced silencing complex (miRISC) that represses target genes containing sequence complementarity to the miRISC-loaded miRNA (Ambros, 2023; Huberdeau and Simard, 2019).

In the nematode *Caenorhabditis elegans*, heterochronic miRNAs serve as molecular switches to inactivate target genes in a stage-specific manner (Rougvie and Moss, 2013; Ivanova and Moss, 2024). The lin-4 miRNA is expressed during the first larval stage (L1) and is required to effectively downregulate the transcription factor *lin-14* at the L1-L2 transition (Wightman et al, 1991; Lee et al., 1993; Wightman et al., 1993; Feinbaum and Ambros, 1999; Greene et al., 2023). Similarly, the let-7 miRNA accumulates during the L4 stage and represses *lin-41* to promote the L4-Adult transition (Reinhart et al., 2000; Slack et al., 2000; Ecsedi et al., 2015; Aeschimann et al., 2019). Loss of *lin-4* or *let-7* function results in failed developmental transitions and aberrant reiteration of stage-specific developmental gene expression patterns.

While critical downstream targets of lin-4 and let-7 during larval development have been identified (Lee et al., 1993; Wightman et al., 1993; Reinhart et al., 2000; Slack et al., 2000; Ecsedi et al., 2015), our understanding of how miRNA activity and miRNA levels are regulated to coordinate key developmental transitions remains incomplete.

The activity of miRNAs is highly regulated throughout development, including at the transcriptional and post-transcriptional levels. RNA polymerase II transcribes most primary miRNA (pri-miRNA) transcripts (Lee et al., 2004). The transcription factor LIN-42, the *C. elegans* homolog of Period, regulates the abundance of heterochronic miRNAs by controlling their rate of transcription (Van Wynsbeghe et al., 2014; Perales et al., 2014; McCulloch and Rougvie, 2014). Rhythmic transcription of *lin-4* and *let-7* is driven by the nuclear hormone receptors NHR-23 and NHR-85, which synchronizes heterochronic miRNA activity to developmental timing (Kinney et al., 2023). Following transcription, pri-miRNA transcripts are typically cleaved by the Microprocessor complex to produce a stem-loop-containing precursor miRNA (pre-miRNA) (Denli et al., 2004; Gregory et al., 2004). Dicer subsequently processes the pre-miRNA to remove the loop segment and generate a double-stranded miRNA duplex (Grishok et al., 2001; Hutvágner et al., 2001; Ketting et al., 2001; Knight and Bass, 2001). The mature miRNA guide strand is selectively loaded into Argonaute to form the miRISC, whereas the other passenger strand is discarded (Khvorova et al., 2003; Scwarz et al., 2003). Each step of miRNA biogenesis is subject to various forms of regulation and can lead to changes in the mature miRNA sequence (isomiR) and alternative functions for a given miRNA (Ha and Kim, 2014; Creugny et al., 2018; Treiber et al., 2019; Medley et al., 2021). After miRNA biogenesis is completed, the activity of miRISC can be regulated by factors influencing the efficacy of gene targeting or impacting the stability of the miRISC-loaded miRNA. In *C. elegans*, clearance of embryo-specific miRNAs before larval development is achieved by the E3 ubiquitin ligase EBAX-1/ZSWIM8, which targets Argonaute for proteasomal degradation and results in destabilization of loaded miRNAs (Shi et al., 2020; Donnelly et al., 2022; Stubna et al., 2024). Several factors have been identified that influence the repression efficiency of miRISC, including many different classes of RNA-binding proteins (van Kouwenhove et al., 2011; Kim et al., 2021). However, how RNA-binding proteins connect miRNA-dependent gene regulation to animal development is not fully understood.

In this study, we report the characterization of *lin-67* (previously E02D9.1) as a regulator of heterochronic miRNA activity in *C. elegans*. LIN-67 is a predicted RNA-binding protein that contains a K Homology (KH) RNA-binding domain and is the *C. elegans* ortholog of the Sam68-like splicing factor KHDRBS2. Loss of *lin-67* function restores regular stage-specific gene expression and developmental timing to hypomorphic *lin-4* and *let-7* mutants. Additionally, we show that *lin-67* genetically interacts with the miRNA-specific Argonaute *alg-1*. In addition to adult sterility, we found that loss of *lin-67* resulted in heterochronic phenotypes indicative of defects in cell fate specification, differentiation, and developmental timing of gene expression. LIN-67 is a nuclear protein expressed throughout animal development and partially colocalizes with the neuronal-specific splicing factor UNC-75. Mutating the LIN-67 KH domain abolished its nuclear localization. LIN-67 negatively regulates lin-4 miRNA levels, likely at the level of transcription. While loss of *lin-67* did not affect *let-7* expression levels significantly, we observed that loss of *lin-67* restores normal levels of let-7 to *alg-1* mutants. Interestingly, each of the miRNAs downregulated considerably in *lin-67* mutants was at least partially restored in *lin-67; alg-1* double mutants, suggesting that LIN-67 may coordinate with ALG-1 to regulate miRNA abundance. Our findings identify LIN-67 as a key regulator of miRNA-dependent gene regulation and place LIN-67 within the heterochronic gene network essential for *C. elegans* development.

## Materials and Methods

### Nematode culture and genetic analysis

All *C. elegans* strains used in this study were derived from the wild-type N2 strain and grown under standard conditions at 20°C (Brenner, 1974). Strains were cultured on nematode growth medium (NGM) plates, and *Escherichia coli* OP50 was used as a food source. A complete list of strains used in this study is provided in Table S1. RNAi interference (RNAi) assays were performed by feeding *Escherichia coli* HT115 expressing double-stranded RNA as previously described with the L4440 feeding empty vector used as a negative control (Kamath et al., 2000). Extrachromosomal arrays were integrated as previously described (Mello et al., 1991).

Developmental phenotypes and fluorescent protein localization were scored using either a Zeiss Axioscope A1 or Leica DM6 upright microscope equipped with DIC and epifluorescence. Hypodermal *pcol-19::GFP* expression was visually scored in young adult animals by examining whether GFP was expressed in hypodermal cells (positive) or restricted to seam cells (negative). Precocious expression of *pcol-19::GFP* was quantified by scoring L4-staged animals. To determine seam cell number, we counted the number of scm::GFP foci between the pharynx and anus of individual animals. Additional vulval inductions were used to quantify the number of pseudovulva. For alpha-amanitin treatment, young adults were transferred to a 1.5 mL tube containing 500 μL of OP50 bacteria supplemented with 0.05% Triton X-100 and 20 μg/mL alpha-amanitin (Sigma-Aldrich), or only 0.05% Triton X-100 as a control. Animals were incubated at room temperature for 3 hours with gentle rocking.

### RNA extraction and isolation

Total RNA was extracted from L4-staged animals that were developmentally synchronized by standard alkaline hypochlorite treatment (Porta-de-la-Riva et al., 2012). Animals were pelleted by centrifugation, flash-frozen in liquid nitrogen, and stored at -80°C until ready for use. The worm pellets were thawed and suspended in a final volume of 250 µL RNase-free water. 1 mL of Trizol was added, and the mixture was vortexed for 10 minutes. 212 µL of chloroform was added, vortexed for one minute, and separated by centrifugation at 19000 x g for 10 minutes at 4°C. The aqueous phase was then recovered, and RNA was re-extracted using an equal volume of phenol:chloroform:isoamyl alcohol (25:24:1, VWR). The RNA was precipitated for 2 hours at -20°C using an equal volume of isopropanol. 1.8 µL of Glycoblue (Invitrogen) was used as a co-precipitant. The RNA pellets were washed using 80% (v/v) ethanol and allowed to air-dry on ice for 10 minutes. The dried pellets were dissolved in RNase-free water and stored at - 80°C. RNA concentration was determined using a Nanodrop spectrophotometer.

### Small RNA sequencing and data analysis

Small RNAs were size selected using a 15% denaturing UreaGel (National Diagnostics, Atlanta, GA). Single-stranded 18mer and 26mer oligonucleotides were combined at a final concentration of 5 µM and used as a molecular ladder in an empty lane. Samples were run for 90 minutes at 150V to resolve small RNAs. To visualize small RNAs, gels were soaked in 1XTBE solution containing 1X SYBR Gold stain (Invitrogen) for 10 minutes and visualized using UV light. Small RNAs migrating between the 16mer and 26mer markers were excised from the gel and soaked in 750 µL of TE buffer supplemented with 0.3 M NaCl overnight while shaking at 550 rpm at 7°C. The supernatant was transferred to a 0.45 µm column and centrifuged at 4°C for 10 minutes at 2500 x g. Small RNAs were precipitated for 2 hours at -20°C using an equal volume of isopropanol and a 10% sample volume of 3M NaOAc. 1.5 µL of Glycoblue (Invitrogen) was used as a co-precipitant. The small RNA pellets were washed once using 80% (v/v) ethanol and air-dried on ice for 10 minutes. The dried pellets were dissolved in 11 µL of RNase-free water. Size-selected small RNA samples were used to make libraries using the Nextflex Small RNA-Seq Kit v3 (PerkinElmer) according to manufacturer instructions. Small RNA libraries were sequenced at the Kansas State University Integrated Genomics Facility using an Illumina NextSeq 500 instrument at 75 cycles. Small RNA sequence data were processed and analyzed as previously described (Panzade et al., 2022). The cutadapt tool (Martin, 2011) was used to process and trim the sequencing reads. The 3’ adapter sequences were first removed using the parameters “-a ATCTCGTATGCCGTCTTCTGCTTGX” and demultiplexed using the parameters “e 1 -a file$:barcodes.fasta”, where the “barcodes.fasta” contains a list of barcode sequences in fasta format. The 3’ Illumina primer sequence was then removed from each read using the parameters “-a TGGAATTCTCGGGTGCCAAGGAACTCCAGTCAC”. Each end of the processed sequencing reads contained four-nucleotide randomers that were removed using cutadapt. The final four nucleotides of each sequencing read were trimmed using the parameters “-u -4”. The first four nucleotides were trimmed using the parameters “-u 4 -m17 - M29”, which also removed processed reads that were shorter than 17 or longer than 29 nucleotides. Read quality was examined using the fastqc tool. Reads were aligned to a list of mature *C. elegans* miRNAs using bowtie (Langmead et al., 2009). The following command was used for bowtie alignments: “-x miR -a --best --strata --norc -v 3”, where “miR” is a bowtie build assembled using a fasta file containing all *C. elegans* mature miRNA sequences obtained from miRbase. The samtools tool (Li et al., 2009) was used to generate, sort, and index bam files to generate read count data. Normalization (reads per million) was calculated by multiplying the number of mapped reads for individual miRNA loci by the total number of mapped miRNA reads divided by one million.

### qPCR and normalization

qPCR experiments were performed using the QuantiNova SYBR Green RT-PCR kit (Qiagen) according to manufacturer instructions. 100 ng of total RNA from L4-staged worms was used as a template for all reactions. An Applied Biosystems 7500 Real Time PCR instrument performed all qPCR reactions. A full list of oligonucleotides used in this study is provided in Table S2.

### Statistical analysis

R statistical software was used to calculate all statistics. All statistics are presented as mean ± one standard deviation unless otherwise specified. Except for data presented in volcano plots, all P-values were calculated using two-tailed *t*-tests assuming equal variance among samples. The *P*-values used in volcano plots were calculated using the DESeq2 package (Love et al., 2014).

## Results and Discussion

### *lin-67* negatively regulates the heterochronic miRNAs lin-4 and let-7

In *C. elegans*, the heterochronic lin-4 miRNA is essential for normal developmental timing, and loss of lin-4 leads to aberrant reiteration of L1-stage specific developmental programs (Lee et al., 2003; Wightman et al., 1993). The hypomorphic *lin-4(ma161)* mutation is a single nucleotide substitution that impacts the ability of lin-4 to downregulate its target *lin-14* (Lee et al., 1993; Ha et al., 1996). As a consequence of abnormal developmental timing, *lin-4(ma161)* animals do not properly express the adult-specific *Pcol-19::GFP* reporter in hypodermal and seam cells during adulthood (Figure 1A) (Carlston et al., 2021). A previous screen for genetic suppressors of *lin-4(ma161)* identified factors that restore adult-specific expression of *Pcol-19::GFP* to *lin-4(ma161) mutants* (Carlston et al., 2021). One of the genetic suppressors was *lin-67* (formerly *E02D9.1*), a putative RNA-binding protein that has been shown to functionally interact with the *let-7* family of miRNAs and the miRNA-associated Argonaute ALG-1 in *C. elegans* (Haskell and Zinovyeva, 2021). We found that the depletion of *lin-67* by RNAi corrected the timing of *Pcol-19::GFP* expression in *lin-4(ma161)* mutants, suggesting that *lin-67* negatively regulates *lin-4* (Figure 1A). Consistent with previous findings, *lin-67(RNAi)* also restored adult-specific expression of *Pcol-19::GFP* to *let-7(n2853)* and *alg-1(ma192)* mutants (Haskell and Zinovyeva, 2021) (Figure 1B, Figure S1). To further support *lin-67* as a regulator of heterochronic miRNAs, we utilized available *lin-67* loss of function deletion alleles (Figure 1C). The *lin-67(tm6625)* allele is a 398 base deletion that removes the third exon of *lin-67* and spans into neighboring introns, while the *lin-67(tm5862)* allele is a 262 base deletion starting from the end of the third exon and extending into the 3’ UTR (Figure 1C). Animals that are homozygous for either *lin-67* deletion allele are sterile, and heterozygotes are maintained on the hT2 balancer chromosome. However, our analyses focused primarily on the slightly larger *lin-67(tm6625)* deletion allele. Quantification of *lin-67* levels by qPCR revealed a significant reduction in *lin-67* expression levels in *lin-67(tm6625)* mutants (0.52 ± 0.30 fold, p<0.02) relative to controls (1.00 ± 0.40 fold) (Figure 1D), possibly due to nonsense-mediated decay (Arribere et al., 2020). Next, we crossed the *lin-67(tm6625)* allele into *alg-1(ma192)* mutants and examined *Pcol-19::GFP* expression in young adult animals (Figure 1E). For these experiments, all animals also carried the *lin-31(n1053)* mutation to prevent the vulval bursting phenotype associated with *alg-1(ma192)* (Zinovyeva et al., 2014). While most *alg-1(ma192)* young adult animals do not properly express *Pcol-19::GFP* in hypodermal cells (12.7 ± 5.4% hypodermal expression), adult-specific hypodermal *Pcol-19::GFP* expression is strongly restored in *lin-67(tm6625); alg-1(ma192)* double mutants (95.5 ± 4.6% hypodermal expression) (Figure 1F), presumably due to increased heterochronic miRNA activity. These data support the hypothesis that *lin-67* negatively regulates heterochronic miRNA activity during *C. elegans* development.

**Figure 1:**
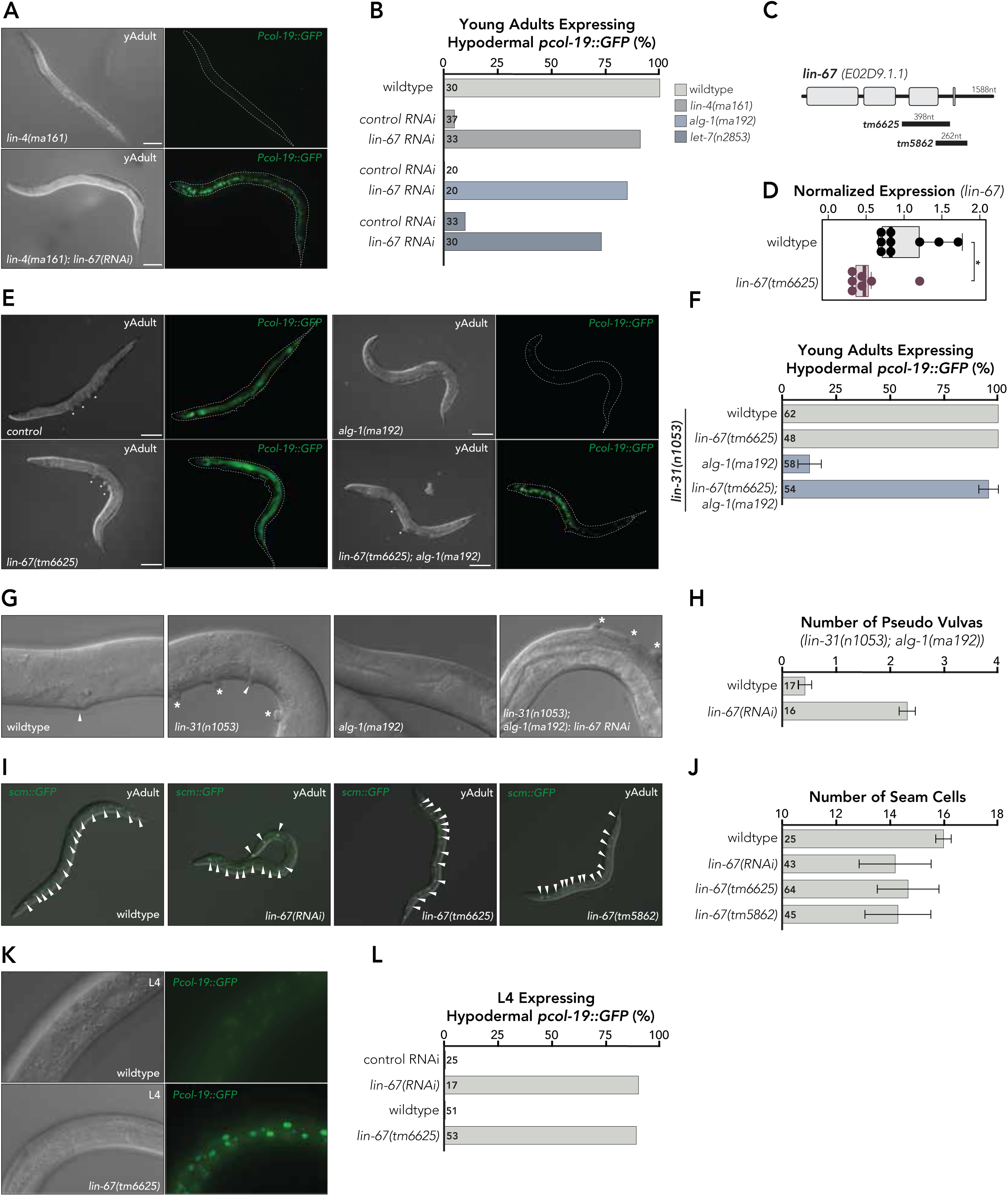
LIN-67 regulates developmental timing. (A) Images of *lin-4(ma161)* young adults expressing the adult-specific *Pcol-19::GFP* reporter. While animals fed control RNAi (L4440) do not properly express hypodermal *Pcol-19::GFP,* depletion of LIN-67 by RNAi restores hypodermal expression of *Pcol-19::GFP* to *lin-4(ma161)* young adults. The scale bar indicates 100µM. (B) Quantification of hypodermal *Pcol-19::GFP* expression in *lin-4(ma161), let-7(n2853)* or *alg-1(ma192)* mutant young adults fed control RNAi or *lin-67(RNAi)*. The number of animals scored (n) for each condition is given to the right of the y-axis. (C) Schematic of *lin-67* gene structure where boxes indicate positions of exons and solid line indicates introns. Solid black bars illustrate the locations of deletion alleles used in this study. (D) Quantification of *lin-67* expression levels by qPCR. Boxplots extend from the first through the third quartile of the data. The solid horizontal line extends 1.5 times the interquartile range or to the minimum and maximum data points. Data were normalized using *tba-1* levels in wild-type animals. *p<0.05 (E) Images of *alg-1(ma192), lin-67(tm6625) and lin-67(tm6625); alg-1(ma192)* mutant young adults expressing the *Pcol-19::GFP* reporter. The *lin-67(tm6625)* mutation restores adult-specific expression to *alg-1(ma192*) mutants. Each strain also carries the *lin-31(n1053)* mutation to suppress the vulval bursting phenotype of *alg-1(ma192)*. Note that *lin-31(n1053)* mutants have a multi-vulva phenotype, but *lin-31(n1053); alg-1(ma192)* double mutants are vulva-less. An asterisk marks vulval structures. The scale bar indicates 100µM. (F) Quantification of hypodermal *Pcol-19::GFP* expression in *alg-1(ma192), lin-67(tm6625) and lin-67(tm6625); alg-1(ma192)* mutant young adults. The number of animals scored (n) for each condition is given to the right of the y-axis. Error bars represent mean ± one standard deviation. (G) Images illustrating vulva and pseudovulva formation in *alg-1(ma192), lin-67(tm6625) and lin-67(tm6625); alg-1(ma192)* mutant young adults that also carry the *lin-31(n1053)* mutation. Arrowheads indicate vulvae, while asterisks indicate pseudovulva formation. alg-1 *(ma192)* mutants are typically vulvaless in the *lin-31(n1053)* genetic background. (H) Quantification of pseudovulva formation. The number of animals scored (n) for each condition is given to the right of the y-axis. Error bars represent mean ± one standard deviation. (I) Images of scm::GFP expression in young adults with reduced *lin-67* function by *lin-67(RNAi)* or the loss of function mutations *lin-67(tm6625)* and *lin-67(tm5862)* illustrate that loss of LIN-67 function results in a reduced number of seam cells. Arrowheads indicate scm::GFP positive seam cells. (J) Quantification of seam cell number. The number of animals scored (n) for each condition is given to the right of the y-axis. Error bars represent mean ± one standard deviation. (K) Images of *lin-67(RNAi)* and *lin-67(tm6625)* L4-stage animals expressing *Pcol-19::GFP*. Note that while hypodermal *Pcol-19::GFP* expression is not observed during the L4 stage in control RNAi or wildtype, *lin-67(RNAi)* and *lin-67(tm6625)* L4-stage animals precociously express hypodermal *Pcol-19::GFP*. (L) Quantification of hypodermal *Pcol-19::GFP* expression in *lin-67(RNAi)* and *lin-67(tm6625)* mutant L4-stage animals. The number of animals scored (n) for each condition is given to the right of the y-axis.

### Loss of *lin-67* results in heterochronic phenotypes

To further understand how *lin-67* regulates animal development, we examined whether the loss of *lin-67* function resulted in additional developmental phenotypes. In *C. elegans*, vulval development begins at the L3 stage when a specific set of precursor cells (P5.p-P7.p) receive an induction signal from the anchor cell (Horvitz and Sternberg, 1991). In *lin-31(n1053)* mutants, the timing of vulval induction is normal. Still, the cellular signals from the anchor cells are amplified, resulting in an over-induction of precursor cells and pseudovulvae formation (Miller et al., 1993) (Figure 1G). In *lin-31(n1053); alg-1(ma192)* double mutants, vulva formation fails due to abnormal timing and amplitude of signals originating from the anchor cell, resulting in a vulva-less phenotype (Figure 1G) (Zinovyeva et al., 2014). We found that RNAi-based depletion of *lin-67* in *lin-31(n1053); alg-1(ma192)* double mutants resulted in a four-fold increase in pseudovulvae formation compared to control RNAi (Figure 1H), suggesting *that lin-67(RNAi)* corrects the induction timing defect of *lin-31(n1053); alg-1(ma192)* double mutants. We also found that loss of *lin-67* function in an otherwise wild-type genetic background resulted in a protruding vulva phenotype (Figure S2), further supporting that *lin-67* regulates vulva development. While we observed 100% adult sterility in homozygous *lin-67(tm6625)* mutants (100% sterile, n=63), we did not observe any restored fecundity in *lin-67(tm6625); alg-1(ma192)* double mutants (100% sterile, n=65), which might suggest that the sterility observed in *lin-67* mutants is unlikely due to hyperactive *alg-1* activity.

Seem cells in the *C. elegans* hypodermis are epithelial cells organized in a singular line across the anterior/posterior axis (Page and Johnstone, 2007). Seam cells complete one round of asymmetric cell divisions during each larval molt, resulting in 16 seam cells by the end of the L4 stage (Gleason and Eisenmann, 2010; Ren and Zhang, 2010). After seam cell proliferation is completed, 16 seam cells fuse and terminally differentiate to form a ridge-like cuticular structure referred to as alae (Page and Johnstone, 2007). The proliferation of seam cells and their terminal differentiation are regulated by heterochronic miRNAs (Resnick et al., 2010; Ambros, 2011). We observed that loss of *lin-67* function resulted in gapped alae (Figure S3), a phenotype that results when animals lack individual seam cells (Page and Johnstone, 2007, Singh and Sulston, 1978). To address whether *lin-67* regulates seam cell proliferation, we used the *scm::GFP* reporter (Terns et al.,1997; Koh and Rothman, 2001) to visualize and count the number of seam cells in young adult *lin-67(RNAi)* and *lin-67* mutant animals (Figure 1I). Compared to wildtype controls that typically contained 16 seam cells (15.97 ± 0.29 seam cells), we observed a reduced number of seam cells in *lin-67(RNAi)* (14.18 ± 1.33 seam cells) or *lin-67(tm6625)* (14.66 ± 1.14 seam cells) and *lin-67(tm5862)* (14.28 ± 1.22 seam cells) mutant animals (Figure 1J). As loss of *lin-67* function restored adult-specific hypodermal expression of *Pcol-19::GFP* to heterochronic miRNA mutant backgrounds (Figure 1A-E), we next examined whether loss of *lin-67* affected the timing of *Pcol-19::GFP* expression throughout animal development (Figure 1K). Whereas hypodermal expression of *Pcol-19::GFP* begins during the late L4 stage in wildtype animals, we found that loss of *lin-67* function resulted in precocious hypodermal expression of *Pcol-19::GFP* during the early L4 stage (Figure 1L). These results suggest that *lin-67* is required for proper cell fate specification and differentiation within the hypodermis.

### LIN-67 localizes to nuclei throughout *C. elegans* development

To gain insight into how *lin-67* regulates developmental timing, we examined the localization of LIN-67 throughout animal development. The *lin-67* gene encodes a predicted RNA-binding protein that contains a conserved K-Homology (KH) RNA-binding domain (Siomi et al., 1993). The closest human homolog of LIN-67 is KHDRBS2/SLM1 (Figure S4), a Sam68-like factor that regulates alternative splicing (Fruscio et al., 1999; Wang et al., 2002; Lijima et al., 2014; Traunmüller et al., 2016; Tan et al., 2022). To examine the localization of LIN-67, we constructed a strain expressing an N-terminal GFP::LIN-67 fusion protein under the control of the native *lin-67* promoter (*Plin-67::GFP::lin-67*). Live imaging revealed that GFP::LIN-67 is ubiquitously expressed throughout animal development and predominantly localizes to nuclei (Figures 2A-E), including the nuclei of hypodermal and vulval precursor cells (Figure 2C) as well as other cell types, including neurons and intestinal cells (Figures 2D-E). In some cases, we observed enrichment of LIN-67 in sub-nuclear foci (Figure 2C), a similar pattern to what has previously been described for the LIN-67 homolog KHDRBS2/SLM1 in HeLa cells (Chen et al., 1999). Interestingly, the localization of LIN-67 to subnuclear foci appears to require active Pol II elongation, as treatment with α-amanitin to block Pol II transcriptional elongation resulted in a more diffuse distribution of LIN-67 throughout the nucleus, an effect that was not observed in α-amanitin resistant *ama-1(m118)* mutants (Rogalski and Riddle, 1988; Bullerjahn and Riddle, 1988; Rogalski et al., 1988; Bird and Riddle, 1989) (Figure S5). We observed a similar localization pattern of LIN-67 in a strain carrying an extrachromosomal array expressing an N-terminal mCherry::LIN-67 fusion protein under the control of the native *lin-67* promoter (*Plin-67::mCherry::lin-67*) (Figure S6). Given that LIN-67 homologs appear to regulate alternative splicing (Lijima et al., 2014; Traunmüller et al., 2016; Tan et al., 2022), we examined whether LIN-67 colocalized with the neuronal splicing factor UNC-75 (Loria et al., 2003; Kuroyanagi et al., 2013; Norris et al., 2014; Chen et al., 2016; Koterniak et al., 2020). As previously reported, we found that GFP::UNC-75 localized to nuclei within the neurons of the ventral cord and head ganglia (Loria et al., 2003; Kuroyanagi et al., 2013) (Figure 2F), whereas expression of mCherry::LIN-67 was observed in the nuclei in all cell types (Figure 2G). We found that mCherry::LIN-67 partially colocalized with GFP::UNC-75 in both the ventral chord and head ganglia (Figure 2H). However, the colocalization appeared to be more pronounced in the neurons of the ventral chord (Figures 2I-L). Taken together, our observations suggest that LIN-67 likely functions within the nucleus and possibly coordinates with other splicing factors to regulate animal development.

**Figure 2.**
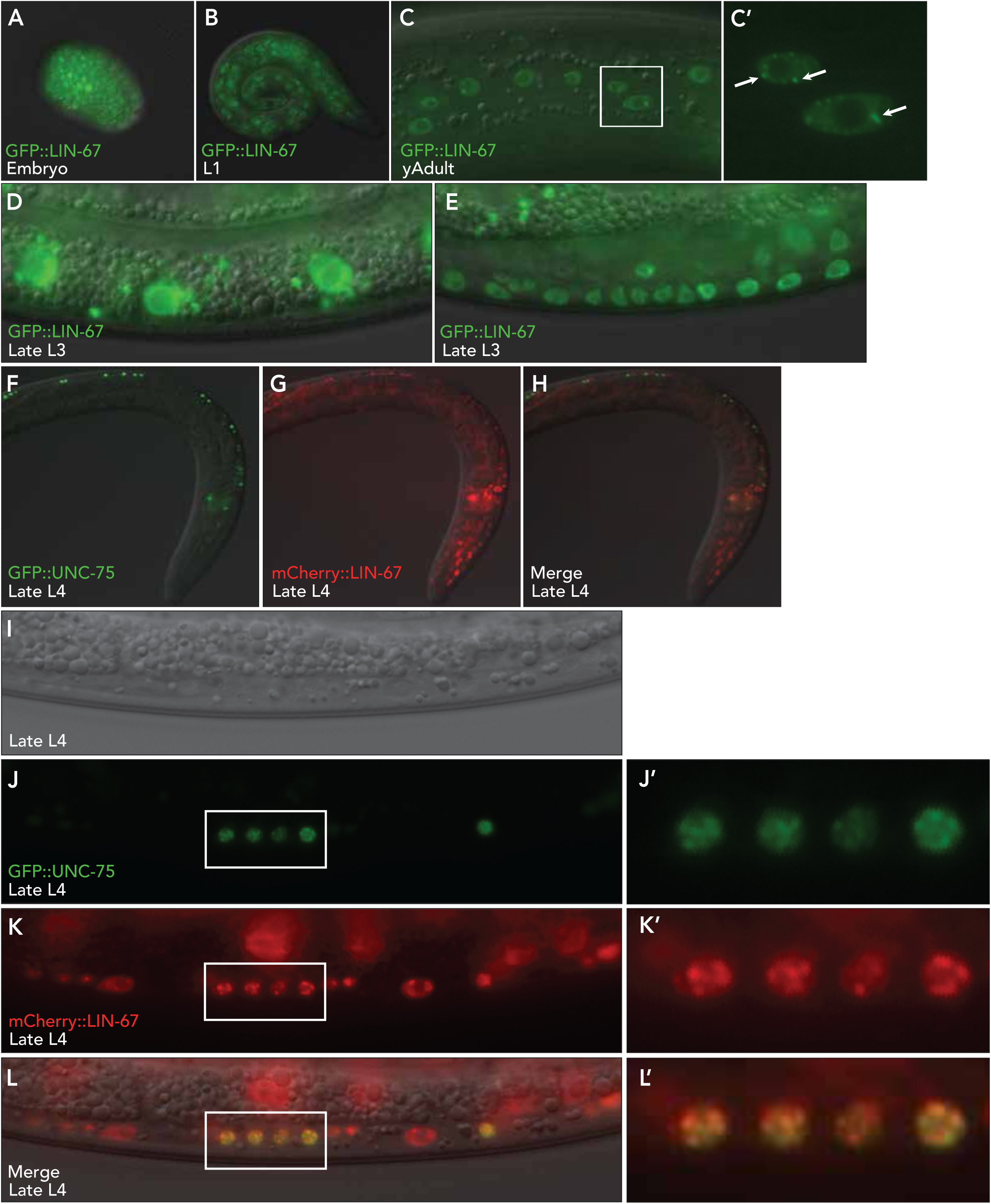
Localization of LIN-67 throughout *C. elegans* development. GFP::LIN-67 localizes to nuclei throughout animal development. Live expression of transgenic Plin-67::GFP::LIN-67 in (A) an embryo, (B) L1-stage, (C) hypodermal cells at young adult, (D) intestinal cells at late L3-stage and (E) vulval precursor cells at late L3-stage (C’) Inset illustrates nuclear localization of GFP::LIN-67 in young adult magnified 3-fold. Arrows indicate enrichment of GFP::LIN-67 in nuclear puncta. (F-H) Transgenic Plin-67::mCherry::LIN-67 promoter partially colocalizes with Punc-75::GFP::UNC-75 in the nuclei of neural cells during the L4 stage. While GFP::UNC-75 is restricted to the ventral cord and head ganglia, mCherry::LIN-67 is expressed in multiple tissues. (I-L) Colocalization of GFP::UNC-75 and mCherry::LIN-67 in motor neurons of the ventral chord. (J’-L’) Insets illustrate the colocalization of UNC-75 and LIN-67 within the nuclei of motor neurons within the ventral chord magnified 3-fold.

### The KH domain of LIN-67 is required for its nuclear localization and animal development

As LIN-67 contains a KH domain that is predicted to bind RNA (Siomi et al., 1993; Valverde et al., 2008; Nicastro et al., 2015), we next asked whether the KH domain of LIN-67 is required for its function during animal development. The interactions between the KH domain and RNA substrate are mediated by a hydrophobic patch that contains a conserved GxxG loop (Hollingworth et al., 2012). Introducing two aspartic acid residues into the GxxG loop (GDDG), abolishes the ability of KH domains to bind RNA without impacting the overall stability of the KH domain (Hollingworth et al., 2012). We first asked whether the LIN-67 KH domain is required for its proper localization throughout animal development. To address this, we generated a strain carrying an extrachromosomal array expressing a mutant mCherry::LIN-67^2D^ (LIN-67^2D^: GDDG, Figure 3A) fusion protein under the control of the native *lin-67* promoter (*Plin-67::mCherry::lin-67^2D^)*. We found that mCherry::LIN-67^2D^ was expressed in a diffuse pattern and failed to localize to the nucleus in various cell types, including hypodermal cells (Figure 3B) and actively dividing vulval precursor cells (Figure 3C). These results support that the KH domain of LIN-67 is required for its nuclear localization, which may be mediated via the interaction of LIN-67 with RNA.

**Figure 3.**
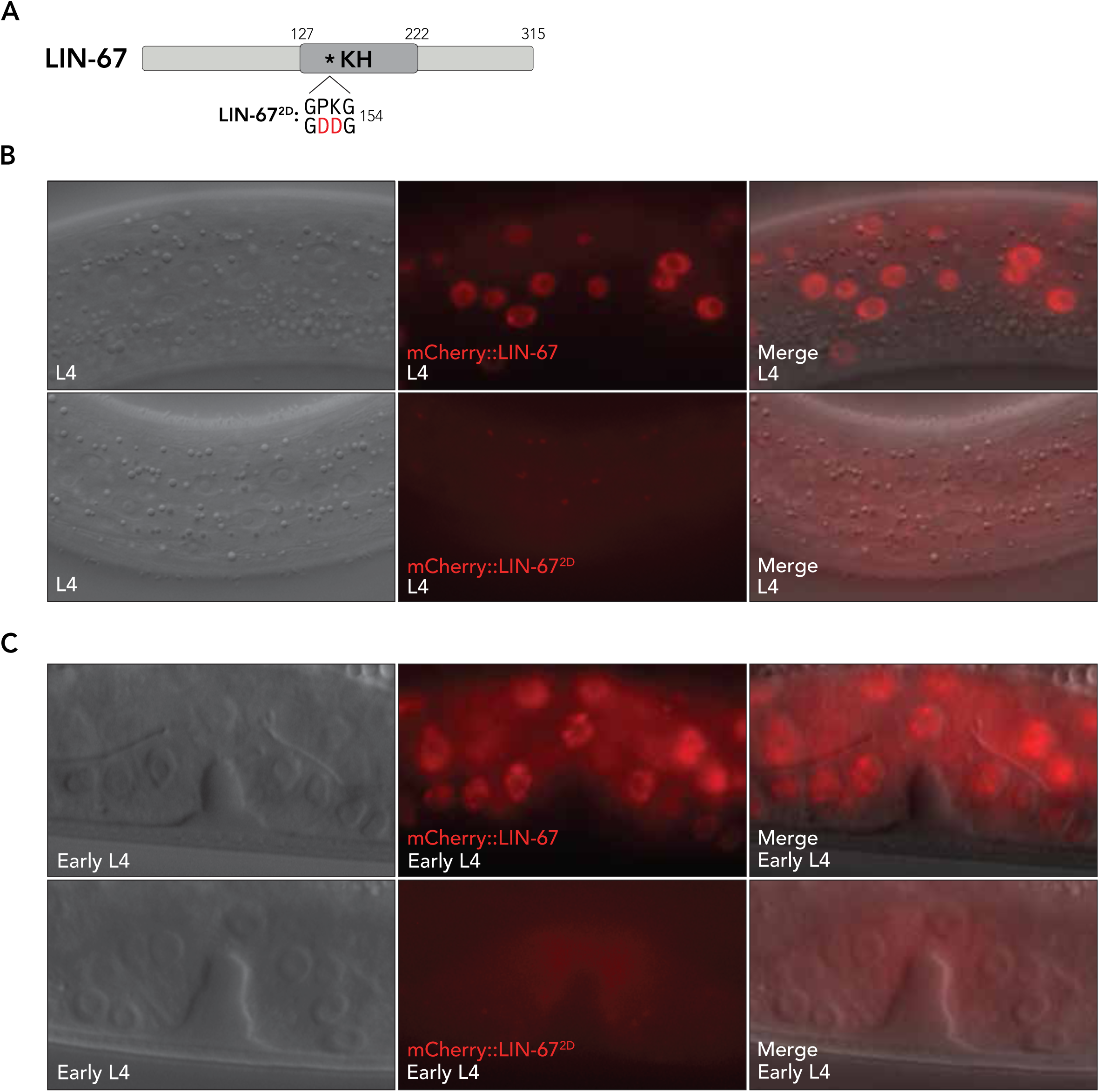
The LIN-67 KH domain is required for its nuclear localization. (A) Schematic representation of predicted LIN-67 functional domains. The asterisk indicates the position of the KH domain mutation (red) used in this study. The 2-nucleotide substitution is predicted to prevent KH-dependent RNA-binding activity of LIN-67. (B) Expression of Plin-67::mCherry::LIN-67 and Plin-67::mCherry::LIN-67^2D^ in L4-stage hypodermal cells. (C) Expression of Plin-67::mCherry::LIN-67 and Plin-67::mCherry::LIN-67^2D^ in L4-stage vulval precursor cells.

### LIN-67 and ALG-1 have opposing effects on miRNA populations

Given our observation that *lin-67* genetically interacts with *alg-1* and heterochronic miRNAs (Figure 1), we next asked whether LIN-67 might regulate the levels of individual miRNAs. To address this, we performed small RNA sequencing in *lin-67* mutant L4-staged animals (Table S3). We found that miRNA populations were broadly disrupted in *lin-67(tm6625)* mutants compared to controls (Figure 4A). Differential expression analysis (DESeq) revealed that a total of 48 miRNAs were significantly (p_adj_<0.01) changed in *lin-67(tm6625)* mutants compared to controls, including 33 downregulated miRNAs and 15 upregulated miRNAs (Figure 4A, Table S3). Interestingly, most downregulated miRNAs in *lin-67(tm6625)* were miRNA passenger strands (30/33 downregulated passenger strands), whereas more of the upregulated miRNAs were guide strands (9/15 upregulated guide strands). These results suggest that loss of *lin-67* may influence passenger strand disposal or stability. Given that miRNA passenger strands are increased in *alg-1(ma192)* mutants (Zinovyeva et al., 2015) and that loss of *lin-67* restores developmental gene expression patterns to *alg-1(ma192)* mutants, we asked how the loss of *lin-67* might affect miRNA populations in the *alg-1(ma192)* mutant background. Consistent with previous findings (Zinovyeva et al., 2015), we found that the levels of some miRNAs were significantly affected in *alg-1(ma192)* mutants compared to wild-type controls (Figure 4B, Table S3). Although several miRNAs were significantly affected in *lin-67(tm6625); alg-1(ma192)* double mutants compared to wildtype, there was only partial overlap in the miRNAs that were significantly affected in *lin-67(tm6625); alg-1(ma192)* double mutants compared to either single mutant (Figure 4C, Table S3). We then examined whether miRNA levels were changed in *lin-67(tm6625); alg-1(ma192)* double mutants compared to *lin-67(tm6625)* or *alg-1(ma192)* single mutants. A few miRNAs, including miR-81 and miR-788 that were significantly changed in *alg-1(ma192)* mutants compared to wildtype showed the opposite change in *lin-67(tm6625); alg-1(ma192)* double mutants (Figure 4D, Table S3), which might at least partially contribute to how *lin-67* negatively regulates *alg-1*. On the other hand, when we compared the miRNA populations of *lin-67(tm6625)* single mutants compared to *lin-67(tm6625); alg-1(ma192)* double mutants, we observed a much higher degree of overlap (Figure 4E, Table S3). Interestingly, all miRNAs that were significantly downregulated in *lin-67(tm6625)* single mutants were comparatively upregulated in *lin-67(tm6625); alg-1(ma192)* double mutants, many of which were restored to near-wildtype levels (Figure 4F). The miRNAs that were upregulated in *lin-67(tm6625)* single mutants compared to wildtype showed little change in *lin-67(tm6625); alg-1(ma192)* double mutants (Figure 4F), suggesting that *alg-1(ma192)* specifically affects the miRNAs downregulated in *lin-67(tm6625)*. As many of the miRNAs affected in *lin-67(tm6625)* mutants were miRNA passenger strands, we next quantified the number of miRNA guide and passenger strands that were at least 2-fold decreased (Figure 4H) or 2-fold increased (Figure 4I) in each mutant background. Consistently, we found that *lin-67(tm6625)* mutants showed a high number of decreased miRNA passenger strands (Figure 4H), which was reduced in *lin-67(tm6625); alg-1(ma192)* double mutants. Similarly, *alg-1(ma192)* mutants showed a high number of increased passenger strands (Figure 4I), which was also reduced in *lin-67(tm6625); alg-1(ma192)* double mutants. Thus, *lin-67* and *alg-1* appear to have countering effects on miRNA populations, especially miRNA passenger strands. As *alg-1(ma192)* partially restores proper miRNA levels to *lin-67(tm6625)* mutants, to some extent, these data support that *alg-1* also acts as a negative regulator of *lin-67*. Our findings suggest that *lin-67* opposes *alg-1* in regulating miRNA abundance, possibly at the level of miRNA biogenesis or in guiding strand selection.

**Figure 4.**
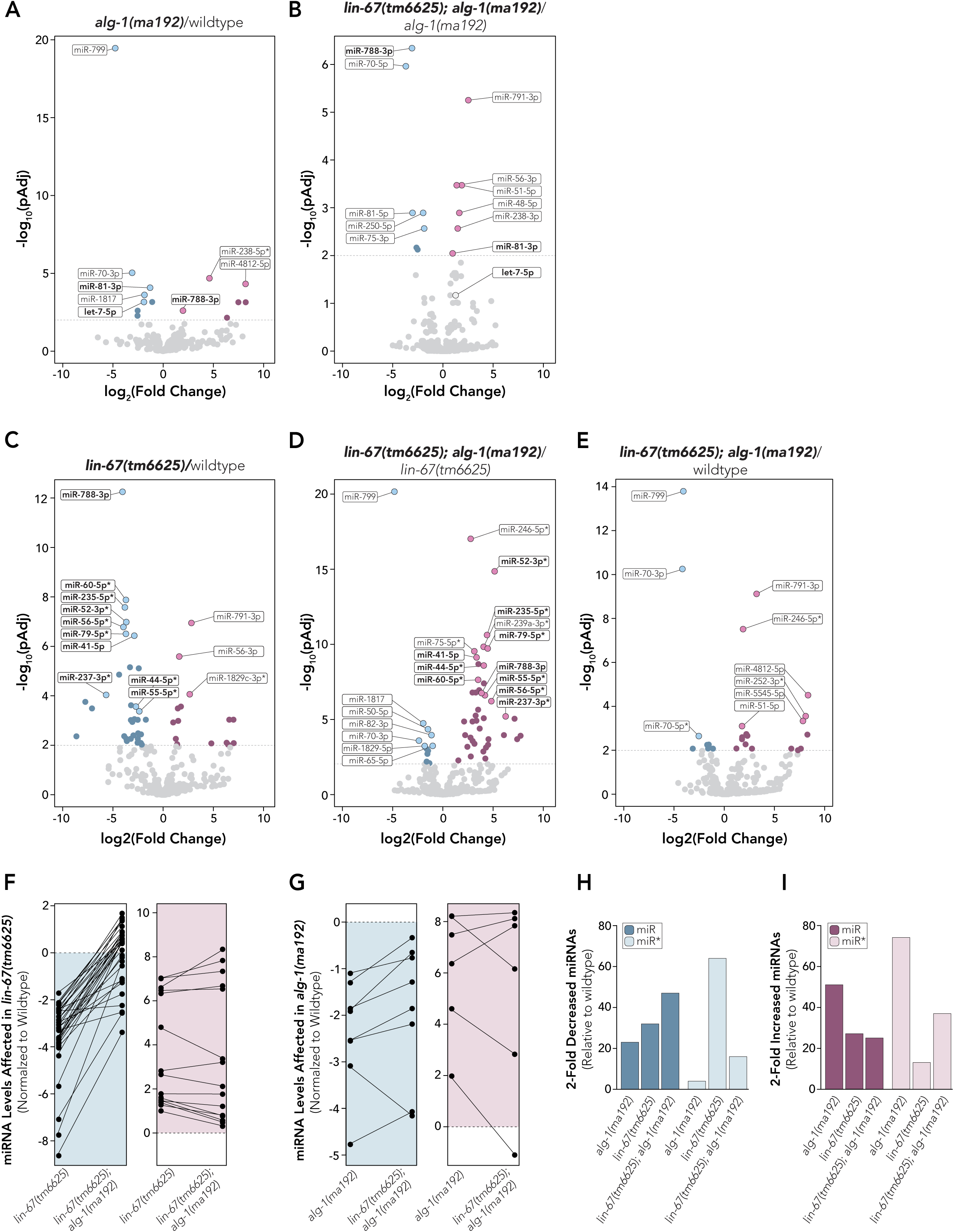
LIN-67 and ALG-1 have opposing effects on total miRNA populations. (A-E) Volcano plots illustrate differences in miRNA populations under different conditions. P-values were calculated using the DESeq2 R package. An adjusted P-value of 0.01 was used as a cutoff to determine statistical significance. Blue dots indicate significantly reduced miRNAs in each sample, whereas purple dots indicate significantly increased miRNAs. Names of miRNAs that are bolded represent miRNAs that were decreased under one condition but increased under a different condition or vice versa. The dashed line represents the significance threshold of p<0.01 (A) Comparison of miRNA populations in *alg-1(ma192)* single mutants to wildtype. (B) Comparison of miRNA populations in *lin-67(tm6625); alg-1(ma192)* double mutants to *alg-1(ma192)* single mutants. (C) Comparison of miRNA populations in *lin-67(tm6625)* single mutants to wildtype. (D) Comparison of miRNA populations in *lin-67(tm6625); alg-1(ma192)* double mutants to *lin-67(tm6625)* single mutants (E) Comparison of miRNA populations in *lin-67(tm6625); alg-1(ma192)* double mutants to wildtype. (F) Changes in the levels of miRNAs that were significantly decreased (left, blue) or increased (right, purple) in *lin-67(tm6625)* single mutants compared to *lin-67(tm6625); alg-1(ma192)* double mutants. Note that many of the miRNAs that were decreased in *lin-67(tm6625)* single mutants compared to wildtype are at least partially restored in *lin-67(tm6625); alg-1(ma192)* double mutants whereas the miRNAs that were increased in *lin-67(tm6625)* single mutants compared to wildtype remain increased in *lin-67(tm6625); alg-1(ma192)* double mutants. (G) Changes in the levels of miRNAs that were significantly decreased (left, blue) or increased (right, purple) in *alg-1(ma192)* single mutants compared to *lin-67(tm6625); alg-1(ma192)* double mutants. (H) Quantification of the number of miRNAs that were decreased at least 2-fold in different mutants compared to wildtype controls. Guide strands (miR) are represented as dark blue, while passenger strands (miR*) are represented by a light blue color (I). Quantification of the number of miRNAs increased at least 2-fold in different mutants compared to wildtype controls. Guide strands (miR) are represented as dark purple, while a light purple color represents passenger strands (miR*)

### LIN-67 regulates the levels of heterochronic miRNAs and influences their activity

Consistent with previous findings (Zinovyeva et al., 2014), we observed a significant decrease in mature let-7 levels in *alg-1(ma192)* mutants (Figure 5A, Table S3). We hypothesized that loss of *lin-67* might restore developmental gene expression to *alg-1(ma192)* mutants by restoring heterochronic miRNA levels. While we did not observe a significant change in mature let-7 levels in *lin-67(tm6625)* single mutants, we found that let-7 levels were significantly increased in *lin-67(tm6625); alg-1(ma192)* double mutants compared to *alg-1(ma192)* single mutants (Figure 5A, Table S3). We also quantified the levels of the let-7 family members miR-48, miR-84 and miR-241, and found that miR-48 levels were significantly increased in *lin-67(tm6625); alg-1(ma192)* double mutants compared to *alg-1(ma192)* single mutants (Figure 5A, Table S3). We did not observe a significant difference in the levels of miR-84 and miR-241 in *lin-67(tm6625); alg-1(ma192)* double mutants compared to *alg-1(ma192)* single mutants, although the levels of miR-241 were significantly decreased compared to wildtype (Figure 5A, Table S3). We also observed a significant increase in mature lin-4 levels in *lin-67(tm6625)* mutants compared to wildtype (Figure 5B, Table S3). However, it is worth noting that our small RNA sequencing analysis was performed using L4-staged animals after the developmental requirement of lin-4 at the L1-L2 transition (Lee et al., 1993). To address how *lin-67* regulates let-7 levels, we examined whether loss of *lin-67* affected primary *let-7* (*pri-let-7*) levels using qPCR. We found that *pri-let-7* levels were significantly increased in *lin-67(tm6625)* (1.96 ± 1.09 fold, p<0.05) and lin-67*(tm6255); alg-1(ma192)* mutants (2.12 ± 0.54 fold, p<0.001) compared to wildtype (1.00 ± 0.25 fold), while pri-let-7 levels were not significantly affected in *alg-1(ma192)* mutants (1.42 ± 0.92 fold, p=0.22) (Figure 5C). Given that loss of *lin-67* does not significantly affect mature let-7 levels, we hypothesize that *lin-67* may negatively regulate *let-7* transcription rather than influencing the rate of pri-let-7 processing. This hypothesis is consistent with our observation that LIN-67 primarily localizes to the nucleus. However, we cannot currently exclude the possibility that LIN-67 also functions downstream of miRNA transcription.

**Figure 5.**
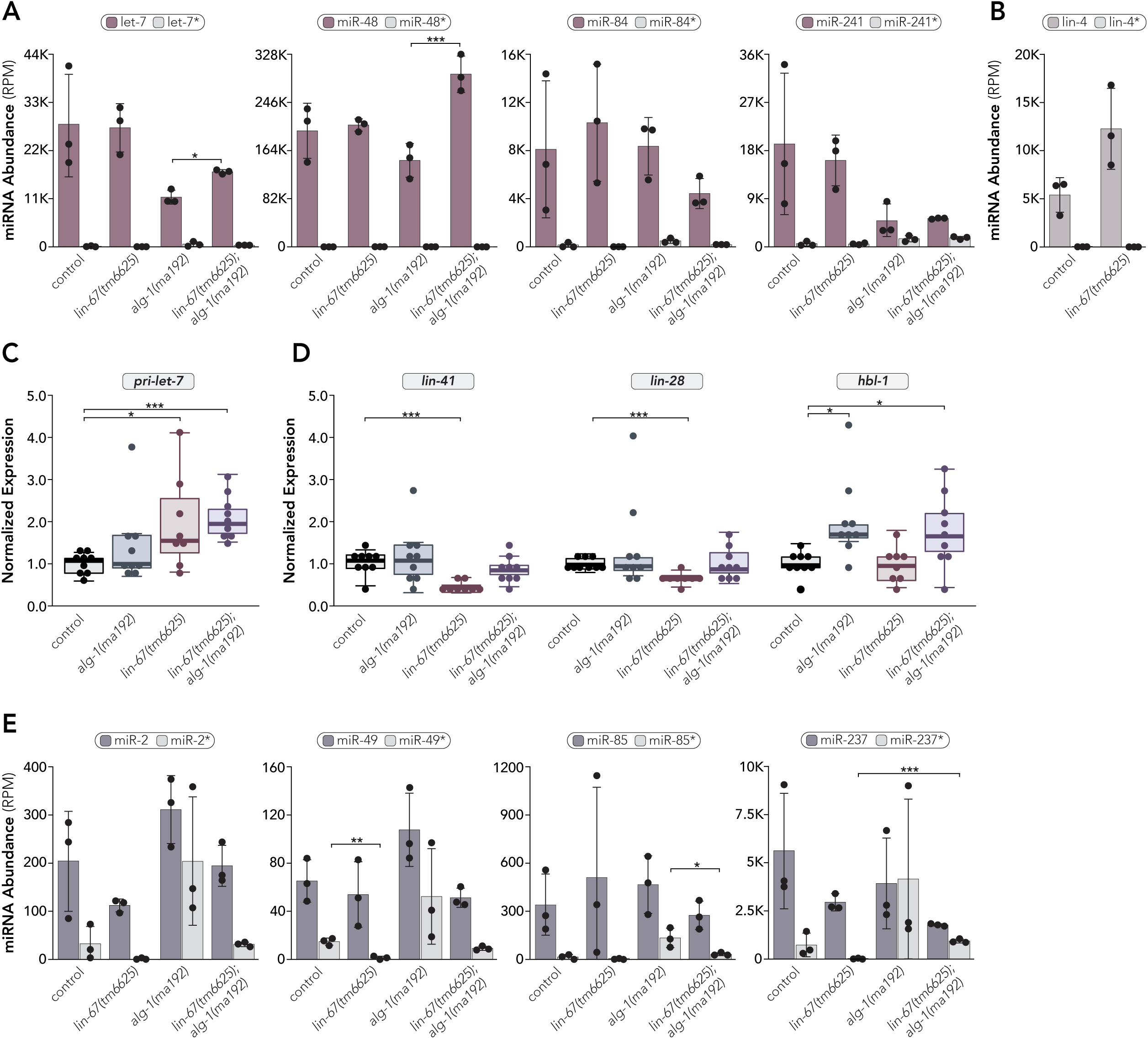
Loss of *lin-67* partially restores levels of heterochronic miRNAs and miRNA passenger strands to *alg-1(ma192)* mutants. (A-C) Normalized read counts (RPM) from small RNA sequencing data. Error bars represent mean ± one standard deviation. Each dot represents one biological replicate. (A) Quantification of the let-7 family. (B) Quantification of lin-4 in *lin-67(tm6625)* mutants compared to wildtype. (C) Quantification of *pri-let-7* expression levels by qPCR. Boxplots extend from the first through the third quartile of the data. The solid horizontal line extends 1.5 times the interquartile range or to the minimum and maximum data points. Data were normalized using *tba-1* levels in wild-type animals. (D) Quantification of expression levels of the heterochronic miRNA target *genes lin-41*, *lin-28* and *hbl-1* by qPCR. Boxplots extend from the first through third quartile of the data. Solid horizontal line extends 1.5 times the inter-quartile range or to the minimum and maximum data point. Data were normalized using *tba-1* levels in wild-type animals. (E) Examples of miRNAs where miR* levels are higher in *alg-1(ma192)* mutants and lower in *lin-67(tm6625)* mutants. (A-E) *p<0.05, **p<0.01, ***p<0.001.

To address whether increased heterochronic miRNA levels led to increased activity, we used qPCR to quantify the expression levels of the let-7 target genes *lin-41* (Reinhardt et al., 2000), *hbl-1* (Abrahante et al., 2003; Lin et al., 2003) and *daf-12* (Großhans et al., 2005). We also quantified the expression levels of *lin-28*, which is regulated by lin-4 and miRNAs belonging to the let-7 miRNA family (Moss et al., 1997; Tsialikas et al., 2017). Consistent with decreased miRNA activity, we found that the expression levels of *hbl-1* and *daf-12* were significantly increased in *alg-1(ma192)* mutants (Figure 5D, Figure S7). We also observed a slight increase in the expression levels of *lin-41* and *lin-28* in *alg-1(ma192)* mutants, although this increase was not statistically significant (Figure 5D). We observed an opposite effect in *lin-67(tm6625)* mutants, where the levels of *lin-41* and *lin-28* were significantly decreased compared to wildtype (Figure 5D), consistent with increased heterochronic miRNA activity. We did not observe any change in the expression levels of *hbl-1* in *lin-67(tm6625)* mutants, and we found that *daf-12* levels were increased (Figure 5D, Figure S7). Thus, *lin-67* appears to influence a subset of heterochronic miRNA targets. Notably, let-7-dependent gene regulation appears to occur in a tissue-dependent fashion (Aeschimann et al., 2019; Lin et al., 2003; Abrahante et al., 2003; Großhans et al., 2005), which might suggest that *lin-67* only regulates heterochronic miRNA activity in certain cell types. In *lin-67(tm6625); alg-1(ma192)* double mutants, we found that the expression levels of *lin-41* and *lin-28* were not significantly different than wildtype, while the expression levels of *hbl-1* and *daf-12* remained significantly increased (Figure 5D, Figure S7). These results suggest that loss of *lin-67* does not restore proper expression levels of all heterochronic miRNA targets to *alg-1(ma192)* and that suppression of *alg-1(ma192)* likely occurs, at least in part, through a subset of heterochronic miRNA targets, possibly in a tissue-specific manner. Given that miRNA passenger strands are increased in *alg-1(ma192)* mutants (Zinovyeva et al., 2015) and decreased in lin-67(tm6625) mutants, we also asked how miRNA passenger strands were affected in *lin-67(tm6625); alg-1(ma192)* double mutants. Although *lin-67* and *alg-1* do not regulate the levels of all miRNA passenger strands (Figure 4), we found several examples where *lin-67* and *alg-1* appear to share common passenger strand targets (Figure 5E). For example, the levels of miR-2* were reduced in *lin-67(tm6625)* mutants and increased in *alg-1(ma192)* mutants, while miR-2* levels were near wildtype in *lin-67(tm6625); alg-1(ma192)* double mutants (Figure 5E). Although it is unclear how *lin-67* might positively regulates miRNA passenger strands, these results suggest that *lin-67* and *alg-1* may oppose each other to regulate the balance of guide strand selection or stability. Collectively, our data support the role of *lin-67* as a regulator of miRNA activity and place lin-67 within the heterochronic pathway that regulates *C. elegans* development.

## Acknowledgements

We want to thank the members of the Hammell and Zinovyeva labs for their helpful comments and discussion of the data and manuscript. We want to thank the Ambros lab for providing some of the strains used in this study. Some strains were provided by the National BioResource Project (NBRP, Tokyo, Japan) and the *Caenorhabditis* Genetics Center (CGC), which is funded by NIH Office of Research Infrastructure Programs (P40 OD010440). We are grateful to Dr. Oliver Hobert for providing the *Punc-75::GFP::unc-75* plasmid.

## Funding

This work was supported by grants from the NIH (R01GM117406 to C.M.H and R35GM124828 to A.Z.). J.C.M was supported by fellowships from the NIH (F32GM148040) and the Kansas INBRE program (P20 GM103418). Additional funds were provided by the Kansas INBRE program (P20 GM103418) and the Johnson Cancer Center at K.S.U.

## Notes

### Competing Interest Statement

The authors have declared no competing interest.

