## Supplemental Data for "LIN-67 functionally interacts with heterochronic miRNAs and regulates developmental timing in *Caenorhabditis elegans*"

**Table S1**: List of *C. elegans* strains used in this study

| Strain | Genotype | Origin |
|---|---|---|
| DR432 | *ama-1(m118) IV* | Sanford et al., 1983 |
| FX5862 | *lin-67(tm5862) I/hT2 (I;III)* | Deletion Mutant Consortium |
| FX6625 | *lin-67(tm6625) I/hT2 (I;III)* | Deletion Mutant Consortium |
| HML2 | *lin-4(ma161) II; maIs105 V [pcol-19::GFP]* | Perales et al. 2014 |
| HML11 | *maIs105 V; lin-2(e1309) let-7(n2853) X* | Perales et al. 2014 |
| HML220 | *unc-119(ed3) III; cshIS21 [plin-67::GFP::LIN-67 + Cbr-unc-119 (+)]* | This study |
| HML243 | *wIs51 V [SCMp::GFP]* | Stec et al. 2020 |
| HML308 | *unc-119(ed3) III; cshEx61 [punc-75::GFP::UNC-75 + plin-67::mCherry::LIN-67 + Cbr-unc-119 (+)]* | This study |
| HML314 | *lin-67(tm5862) I/hT2 (I;III); maIs105 V* | This study |
| HML316 | *unc-119(ed3) III; cshEx62 [punc-75::GFP::UNC-75 + plin-67::mCherry::LIN-67$^{2D}$ + Cbr-unc-119 (+)]* | This study |
| VT2077 | *lin-31(n1053) II; maIs105 V* | Perales et al. 2014 |
| VT2079 | *lin-31(n1053) II; maIs105 V; alg-1(ma192) X* | Zinovyeva et al. 2014 |
| UY405 | *lin-67(tm5862) I/hT2 (I;III); lin-31(n1053) II; maIs105 V* | This study |
| UY411 | *lin-67(tm5862) I/hT2 (I;III); lin-31(n1053) II; maIs105 V; alg-1(ma192) X* | This study |

**Table S2**: List of oligonucleotides used in this study

| Name | Sequence (5'-3') | Info |
|---|---|---|
| lin-67-F1 | ACCTCGGGTGATCAGTTTAGAA | *lin-67* Forward qPCR |
| lin-67-R1 | GAAGTTGAAGTTTGGATGACGG | *lin-67* Reverse qPCR |
| pri-let-7-F1 | AAGCAGGCGATTGGTGGAC | *let-7* Forward qPCR |
| pri-let-7-R1 | GTTGTGAGAGCAAGACGACG | *let-7* Reverse qPCR |
| lin-41-F1 | ACATCCTGGAAAGCATCGAG | *lin-41* Forward qPCR |
| lin-41-R1 | AAGCGTTGACGTGTGTATCG | *lin-41* Reverse qPCR |
| lin-28-F1 | TAAACCATACTACCACCTACCT | *lin-28* Forward qPCR |
| lin-28-R1 | AACAGGTGCAATCAGTTCTAT | *lin-28* Reverse qPCR |
| hbl-1-F1 | CTCGTCTAGTGACCCATTCT | *hbl-1* Forward qPCR |
| hbl-1-R1 | ACGCCCGAACATTGATAAG | *hbl-1* Reverse qPCR |
| daf-12-F1 | GTTTAGAGTTCTTCGGTTTCTTCGACGAGG | *daf-12* Forward qPCR |
| daf-12-R1 | CGTTCATCGGAGGATCAGAGCGGACAGAG | *daf-12* Reverse qPCR |

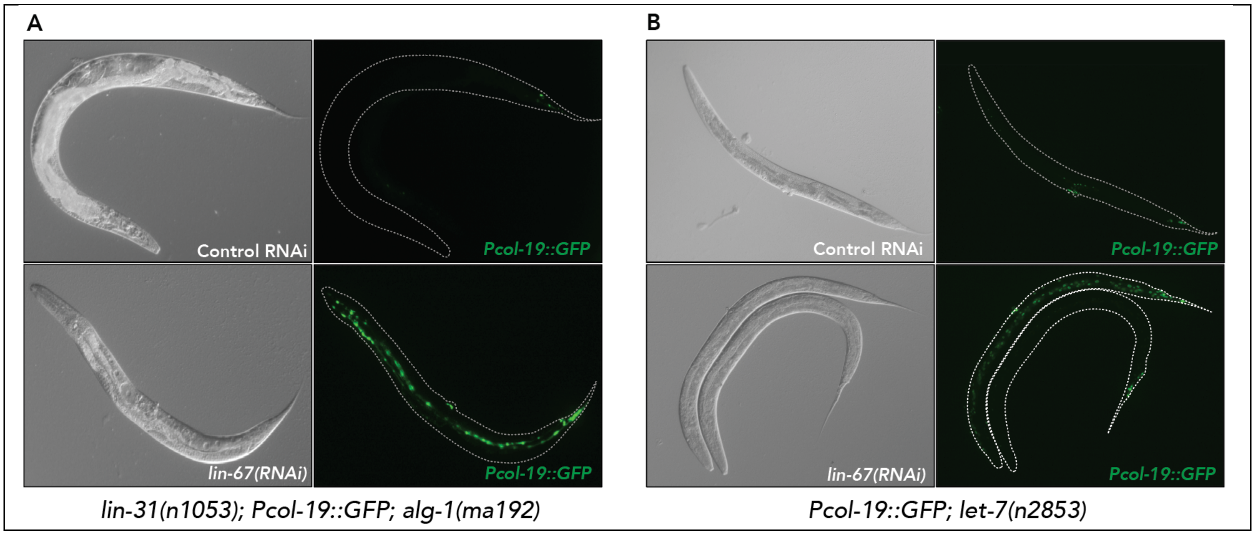


**Figure S1:** *lin-67(RNAi)* restores hypodermal adult-specific *Pcol-19::GFP* expression to (A) *alg-1(ma192)* and (B) *let-7(n2853)* mutants. The L4440 empty RNAi feeding vector was used as a negative control.

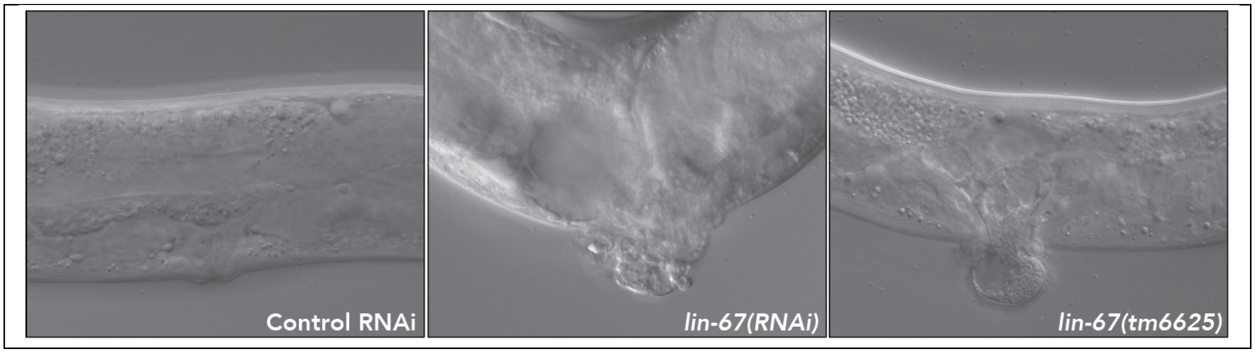


**Figure S2:** Loss of *lin-67* function results in a protruding vulva phenotype. The L4440 empty RNAi feeding vector was used as a negative control.

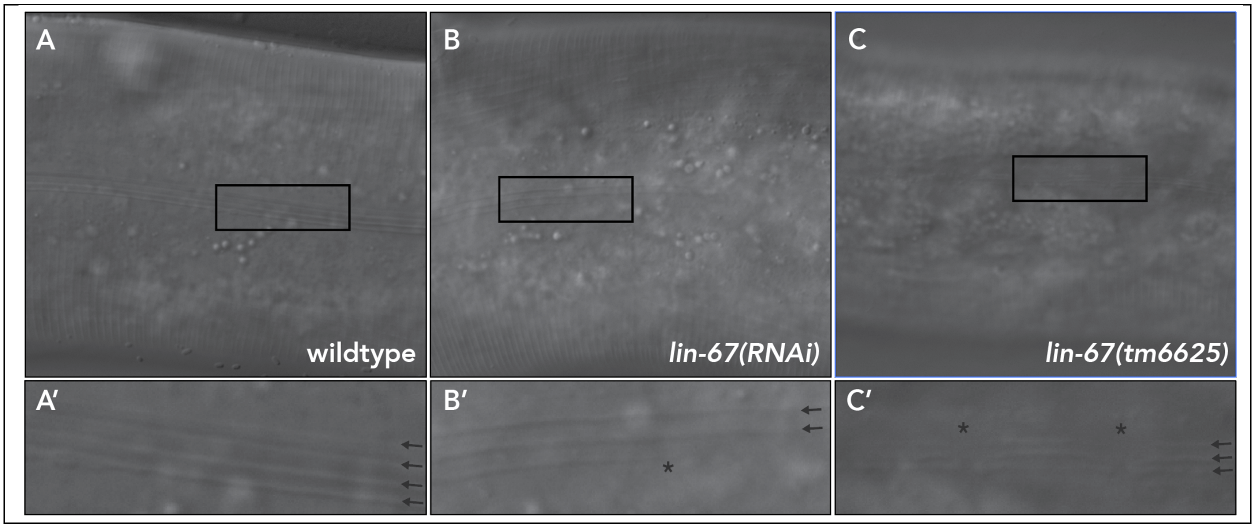


**Figure S3:** Loss of *lin-67* function disrupts alae formation. The L4440 empty RNAi feeding vector was used as a negative control. Insets are magnified 3-fold. Arrows indicate alae tracks and asterisks illustrate missing or gapped alae.

```
LIN-67       MDPYQQGGRGGGFPARGGRGGGHGGGYPQEGYGAAAGGYGGYDPYNPYGAAGGYGMYPGQ   60
KHDRBS2      -------------------------------------------------------MEEE   4
                                                                        :

LIN-67       GYPPQEMTSPLDAEIQAVLREIHLEVTGLETSGDQFRNARRLLNSEKERL------ENNI   114
KHDRBS2      KYLP----------------ELMAEK---DSLDPSFVHASRLLAEEIEKFQGSDGKKEDE   45
              * *                *:  *    :: . .* :* *** .* *::      :::

LIN-67       DPEWLEVDVAKPVKVCKKILVPIYRHPNFNFIGKVLGPKGATLQTLCKTHKCHIYILGRG   174
KHDRBS2      EKKYLDVISNKNIKLSERVLIPVKQYPKFNFVGKLLGPRGNSLKRLQEETGAKMSILGKG   105
             : ::*:*   * :*:.:::*:*: ::*:***:**:***:* :*: * :   .:: ***:*

LIN-67       STKDREKEAELLASGDPQHAHFSGPLHVKVETVAPAYIAYGRVAAVIEELSRILQPIHED   234
KHDRBS2      SMRDKAKEEELRKSGEAKYAHLSDELHVLIEVFAPPGEAYSRMSHALEEIKKFLVPDYND   165
             * :*: ** **  **: ::**:*. *** :*..**   **.*:: .:**:.::* * ::*

LIN-67       TTPAHLKNGSGDGEEKEDEEKKEGGEGGGRGGRGGRGGFRGGFRGGRGGFGGPGGPMGGR   294
KHDRBS2      EIRQEQLRELSYL-----NGSEDSGRGRGIRGRGIRIAPTAPSR-GRGGAIPP-PPPPGR   218
                 .  .  .       :  .::.*.* *  *** * .  .  * ****   *  *  **

LIN-67       GGMGGMGGRGRGG--------------------QAAGFRPY-------------------   315
KHDRBS2      GVLTPRGSTVTRGALPVPPVARGVPTPRARGAPTVPGYRAPPPPAHEAYEEYGYDDGYGG   278
             * :   *.    *                     . *:*

LIN-67       ------------------------------------------------------------   315
KHDRBS2      EYDDQTYETYDNSYATQTQSVPEYYDYGHGVSEDAYDSYAPEEWATTRSSLKAPPQRSAR   338


LIN-67       -----------   315
KHDRBS2      GGYREHPYGRY   349
```

**Figure S4:** Alignment of *C. elegans* LIN-67 and human KHDRBS2/SLM1 amino acid sequences. Identical residues are marked by asterisks and chemically similar residues are marked by colons or periods. The KH domain of each protein is highlighted in yellow and the conserved GxxG motif is indicated by bold text.

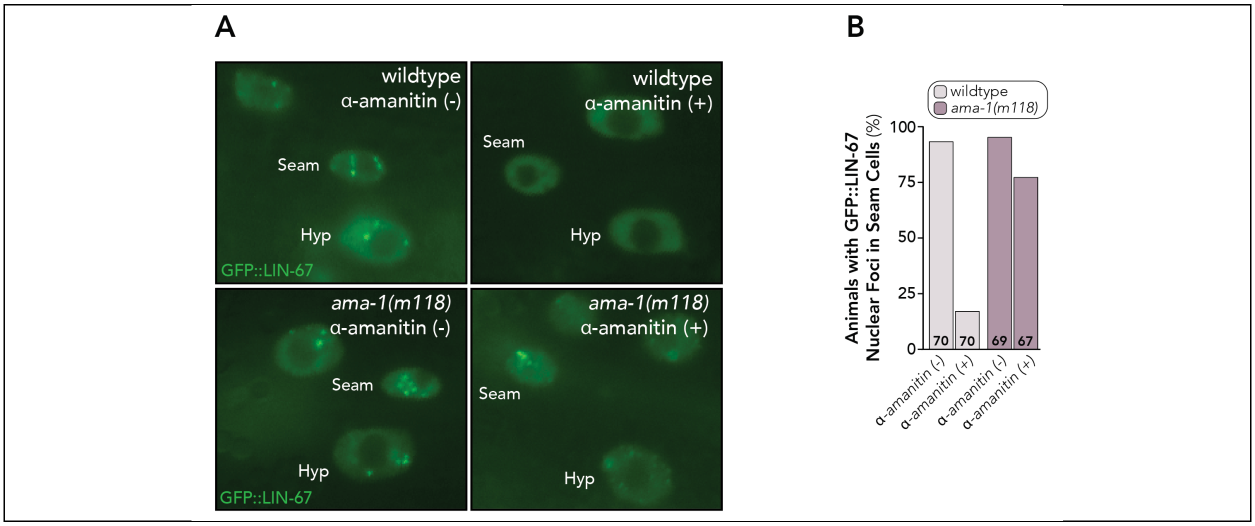


**Figure S5:** α-amanitin disrupts the localization of LIN-67 to subnuclear foci. (A) Subcellular localization of *Plin-67*::GFP::LIN-67 in hypodermal (Hyp) and seam cells. Treatment with α-amanitin (+) disrupts the localization of GFP::LIN-67 to subnuclear foci in wildtype animals, but not in the α-amanitin resistant *ama-1(m118)* mutants. (B) Quantification of the percent of animals that express GFP::LIN-67 within subnuclear foci in seam cells. The number of animals scored (n) is given at the bottom of each bar.

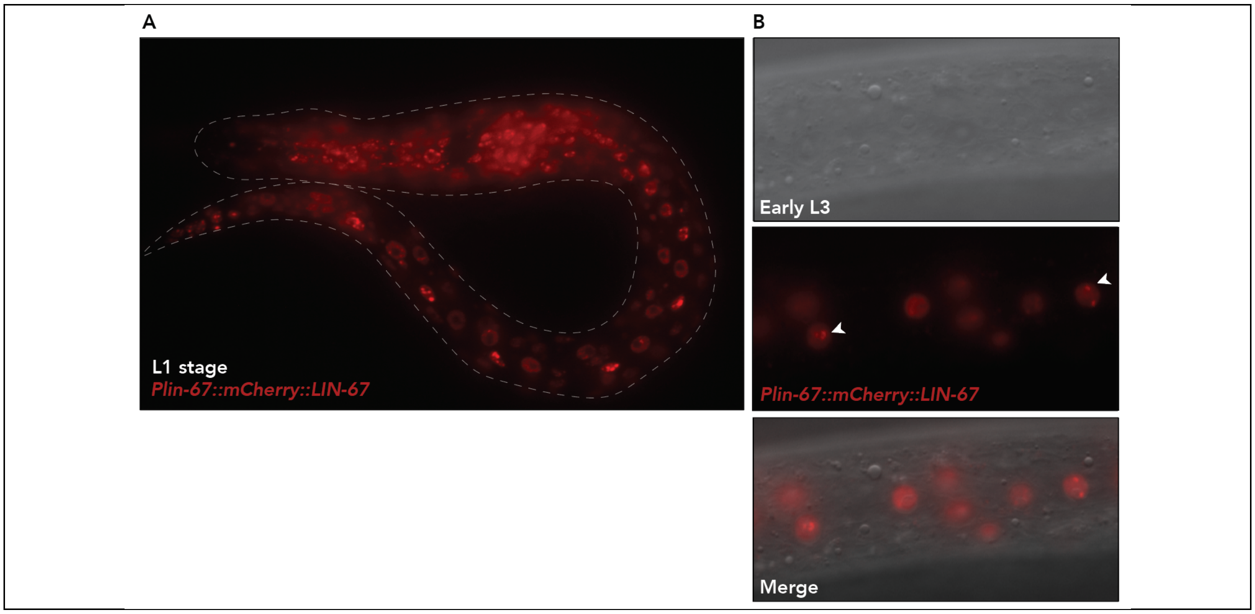


**Figure S6:** Localization of *Plin-67::mCherry::LIN-67* during the (A) L1 stage and (B) hypodermal cells of L3 stage animals. Arrowheads indicate localization of mCherry::LIN-67 to subnuclear foci in the nuclei of hypodermal cells.

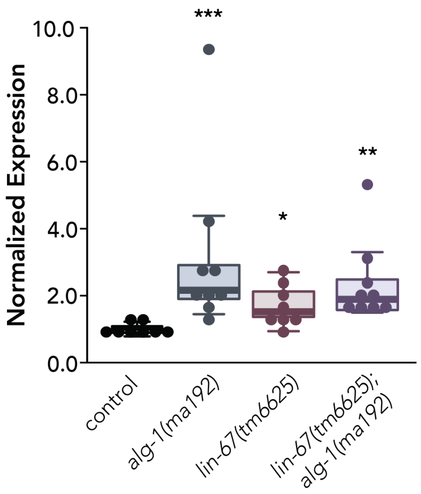


**Figure S7:** Quantification of *daf-12* expression levels by qPCR. Boxplots extend from the first through third quartile of the data. Solid horizontal line extends 1.5 times the inter-quartile range or to the minimum and maximum data point. Data were normalized using *tba-1* levels in wildtype animals. *p<0.05, **p<0.01, ***p<0.001.
